# Antibacterial Activity Potential of Lactic Acid Bacteria (LAB) Isolates from Palm Sap (Arenga pinnata) from the Wawo Plantation, Tomohon City, North Sulawesi

**DOI:** 10.64898/2026.08.22.746455

**Authors:** Yeanly Wuena Pinaria, Noldy Paulus Pangkerego, Geyby Kumolontang

## Abstract

Lactic acid bacteria (LAB) are one of the dominant groups of bacteria in the palm sap (*Arenga pinnata*) microbiome. Previous research in the sago palm sap production centers of Tomohon City (Kayawu, Pinaras, and Lahendong) has successfully identified various LAB species, including *Lactobacillus casei, Lactobacillus plantarum, Lactobacillus brevis, Lactobacillus buchneri, Leuconostoc mesenteroides*, and *Leuconostoc* sp. This study aims to identify LAB species in sago palm sap from a new location, namely the Wawo Plantation in Tomohon, and to evaluate their potential as natural antibacterial agents. Through 16S rDNA gene sequencing analysis, the isolates obtained were identified as belonging to the newly described genera *Lacticaseibacillus* and *Lactiplantibacillus*. Four promising isolates—*Lactiplantibacillus fabifermentans* A1.4, *Lacticaseibacillus casei* B1.5, *Lacticaseibacillus paracasei* B1.6, and *Lacticaseibacillus paracasei* B3.5—were tested for their inhibitory activity against the enteric pathogens *Salmonella* sp. and *Escherichia coli* using the well diffusion method. The results showed that all isolates exhibited a strong spectrum of pathogen inhibition. The highest inhibitory activity against *Salmonella* sp. was demonstrated by the *L. paracasei* B1.6 isolate, with an inhibition zone of 21.25 mm, while optimal inhibition against *E. coli* was achieved by *L. casei* B1.5 at 11.0 mm. These findings confirm that the local BAL strain from Tomohon palm sap has great potential for large-scale development as a biopreservative in the food industry and as a functional probiotic agent.

## 1. Introduction

Palm sap (*Arenga pinnata*) is a non-timber forest product that plays an important economic and cultural role, particularly in North Sulawesi. Palm sap is rich in sucrose, minerals, and vitamins, making it an ideal natural substrate for the growth and proliferation of various microorganisms (Sulistiani et al., 2020). Various food microbiology studies have demonstrated that the group of bacteria that grows dominantly and plays a crucial role in the success of the natural fermentation process of palm sap is the Lactic Acid Bacteria (LAB) group.

The city of Tomohon is known as one of the leading centers for palm sap production in North Sulawesi Province. Several neighborhoods that have historically served as production hubs include Kayawu, Pinaras, and Lahendong, while the Wawo Plantation area has recently begun to be explored on a large scale. The microbial biodiversity of palm sap in the Tomohon region is very high. Several species that have been conventionally isolated and identified from previous studies include *Lactobacillus casei, Lactobacillus plantarum, Lactobacillus brevis, Lactobacillus buchneri, Leuconostoc mesenteroides*, and *Leuconostoc* sp. (Pinaria et al., 2016).

The city of Tomohon is one of the leading centers for palm sap production in North Sulawesi Province. Several neighborhoods that have historically served as production centers include Kayawu, Pinaras, and Lahendong, and the Wawo Plantation area is currently being explored on a large scale. Based on previous studies, the microbial biodiversity of palm sap in the Tomohon region is very high. Several species that have been successfully isolated and identified using conventional methods include *Lactobacillus casei, Lactobacillus plantarum, Lactobacillus brevis, Lactobacillus buchneri, Leuconostoc mesenteroides*, and *Leuconostoc* sp. (Pinaria et al., 2016). In recent developments in molecular taxonomy, the genus *Lactobacillus* has undergone a comprehensive reclassification, in which certain species have been placed into new genera such as *Lactiplantibacillus* (Zheng et al., 2020). The identification of these potential local strains aligns with research conducted by (Pinaria et al., 2023) and (Pinaria & Pangkerego, 2025), which intensively explored the potential of the local *Lactobacillus casei* AL.15 strain from Tomohon palm sap as a biopreservative agent in the food industry. The findings of this study further confirm that the BAL isolate from Tomohon palm sap is capable of secreting bacteriocins and organic acids that are highly effective in inhibiting the growth of pathogens.

An essential function required of BAL isolates—whether as candidate probiotics or commercial biopreservatives—is their ability to produce broad-spectrum antimicrobial compounds to inhibit foodborne *pathogens*. Bacteria such as *Salmonella* sp. and *Escherichia coli* are often the primary agents causing outbreaks of gastrointestinal infections in developing countries. Therefore, this study focused on the isolation and molecular identification process using 16S rDNA sequencing of BAL isolates from the Wawo Plantation. Furthermore, the antibacterial activity of these isolates was tested in vitro against *Salmonella* sp. and *E. coli* to assess their efficacy compared to findings regarding antibacterial profiles in previous studies.

## 2. Research Methodology

### 2.1 Sampling

Fresh palm sap (*Arenga pinnata*) samples were collected directly during the tapping process at the Wawo Plantation in Tomohon City. The sap samples were immediately collected in sterile containers and transported to the laboratory using *a cool box* to maintain a cold temperature (4°C). This cold chain procedure was implemented to prevent cross-contamination and inhibit the rate of spontaneous fermentation by yeast during transport.

### 2.2 Isolation of Lactic Acid Bacteria (LAB)

Bacterial isolation was performed using selective *de Man, Rogosa, and Sharpe Agar* (MRS) medium enriched with 1% CaCO_3_. This modification was intended to easily select and distinguish acid-producing bacteria, which are characterized by the formation of a clear zone (*halo*) around the colonies. Palm sap samples were serially diluted by from^10−1^ to^10−6^, then inoculated onto the surface of the medium using the *spread* plate method. The culture plates were incubated under facultative anaerobic conditions at 37°C for 48 hours. Colonies exhibiting macroscopic morphological characteristics of BAL and clear zone formation were isolated for repeated purification.

### 2.3 Molecular Identification (16S rDNA Gene Sequencing)

Genomic DNA was extracted from selected BAL strains using a commercial kit that meets molecular laboratory standards. Amplification of the 16S rDNA gene fragment was performed via a *Polymerase Chain Reaction* (PCR) using a pair of universal primers, namely Forward 27F and Reverse 1492R. The resulting PCR amplicons were then sequenced. The obtained nucleotide sequence data were analyzed and *aligned* with the *National Center for Biotechnology Information* (NCBI) genetic database using BLAST (*Basic Local Alignment Search Tool*) software to confirm species identity phylogenetically and with precision.

### 2.4 Antibacterial Activity Assay

The antibacterial activity of BAL isolates was evaluated against cultures of the pathogenic bacteria *Salmonella* sp. and *Escherichia coli* using *the well diffusion method*. Test bacterial suspensions with turbidity levels equivalent to the McFarland standard were spread evenly on the surface of *Mueller-Hinton Agar* (MHA). Wells (of standard diameter) were made in the agar, and then *the Cell-Free Supernatant* (CFS) obtained by centrifugation from the cultures of the four BAL isolates was pipetted and added to these wells. The Petri dishes were incubated at 37°C for 24 hours. The antimicrobial inhibitory activity was evaluated by measuring the diameter of the inhibition zone (clear zone) formed around the wells in millimeters (mm).

## 3. Results and Discussion

### 3.1 Identification and Characteristics of BAL Isolates from the Wawo Plantation

Field exploration focused on palm sap samples from the Wawo Plantation in Tomohon successfully yielded a number of pure isolates. Based on the results of basic macroscopic and microscopic biochemical testing, all isolate strains exhibited the essential characteristics of the Lactic Acid Bacteria group. The four leading isolates were Gram-positive (+), had rod-shaped (bacillus) cells, were unable to produce catalase (catalase-negative), and lacked motility structures (non-motile) (Table 1).

**Table 1.** Morphological Characteristics of PAL Isolates from Wawo Palm Sugar Plantation.

| Isolate Code | Gram Stain Test | Cell Morphology | Catalase Test | Motility Test |
| --- | --- | --- | --- | --- |
| A1.4 | + | Batang | - | - |
| B1.5 | + | Batang | - | - |
| B1.6 | + | Batang | - | - |
| B3.5 | + | Batang | - | - |

Further characterization was conducted at the molecular level. The results of 16S rDNA nucleotide sequence homology analysis of the four leading strains confirmed that these isolates occupy taxonomic branches within the groups *Lacticaseibacillus casei, Lacticaseibacillus paracasei*, and *Lactiplantibacillus fabifermentans*, in accordance with the revised genus descriptions (Zheng et al., 2020). The findings in Table 2 conclude that the ecological s of palm sap in the Wawo highlands provide a microenvironment that persistently supports the dominance of vegetative cells of the *L. casei* and *L. paracasei* biotype groups.

**Table 2.** 16S rDNA Sequencing Results for Palm Sap BAL Isolates.

| Isolate Code | Similarity Level | Sequence Length (bp) | Closest Species (NCBI) | Species (Scientific Name) | Accession Number (Reference) |
| --- | --- | --- | --- | --- | --- |
| A1.4 | 99.93% | 1491 | *Lactiplantibacillus fabifermentans* strain DSM 21115 | *Lactiplantibacillus fabifermentans* | NR_113339.1 |
| B1.5 | 99.93% | 1466 | *Lactobacillus casei* strain 067 | *Lacticaseibacillus casei* | JN560867.1 |
| B1.6 | 100.00% | 1483 | *Lactobacillus paracasei* strain 132Y | *Lacticaseibacillus paracasei* | MK774551.1 |
| B3.5 | 100.00% | 1467 | *Lactobacillus paracasei* strain 2738 | *Lacticaseibacillus paracasei* | MT611749.1 |

**Figure 1.**
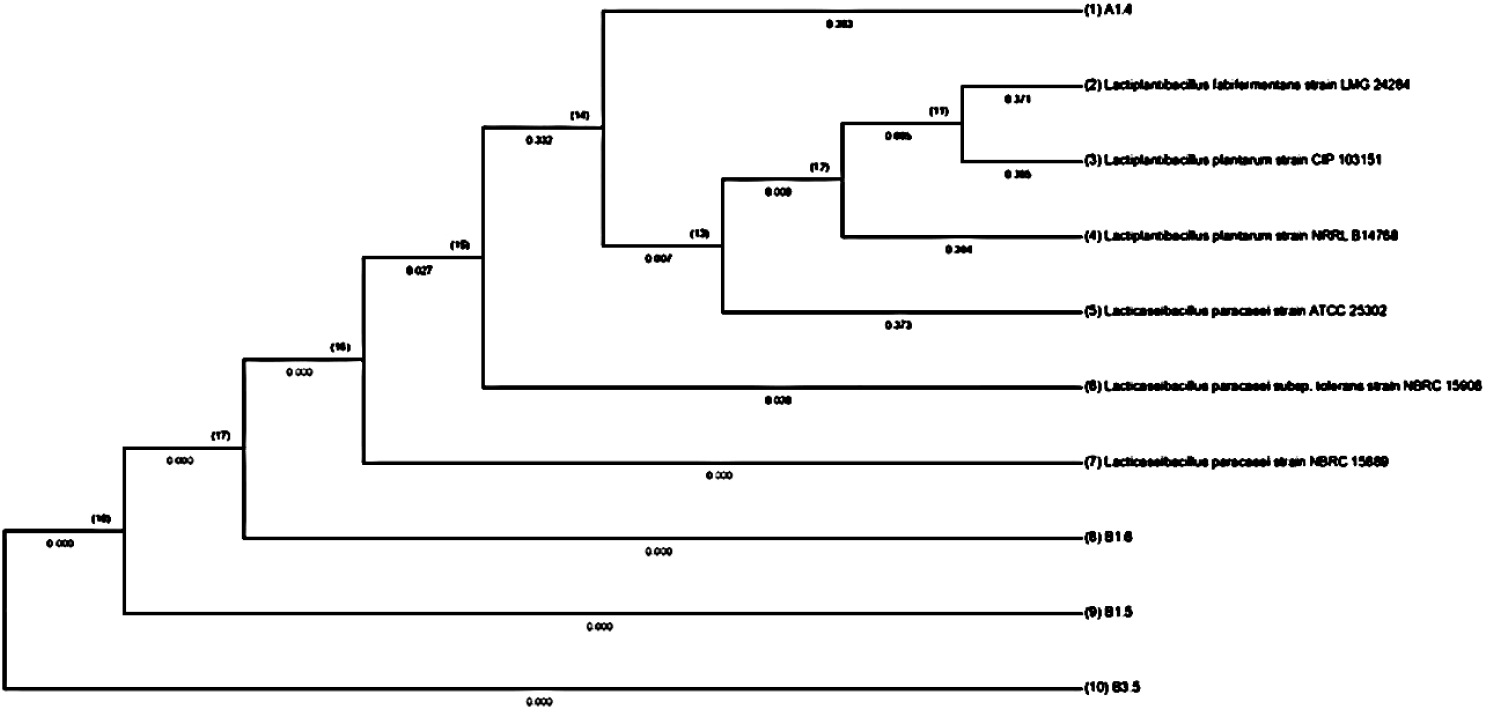
Phylogenetic tree of BAL from Wawo Plantation’s palm sap using Clustalx.

### 3.2 Potential Antibacterial Activity Against Target Pathogens

The ability of these isolates to synthesize and secrete bioactive compounds was evaluated based on the inhibitory activity of their cell-free supernatant (CFS) against test enteropathogenic bacteria. This test followed the standard for measuring inhibition zones in millimeters (mm). Experimental observations revealed very distinct variations in antimicrobial resistance profiles among the four isolates from Wawo, as shown in Table 3.

**Table 3.**
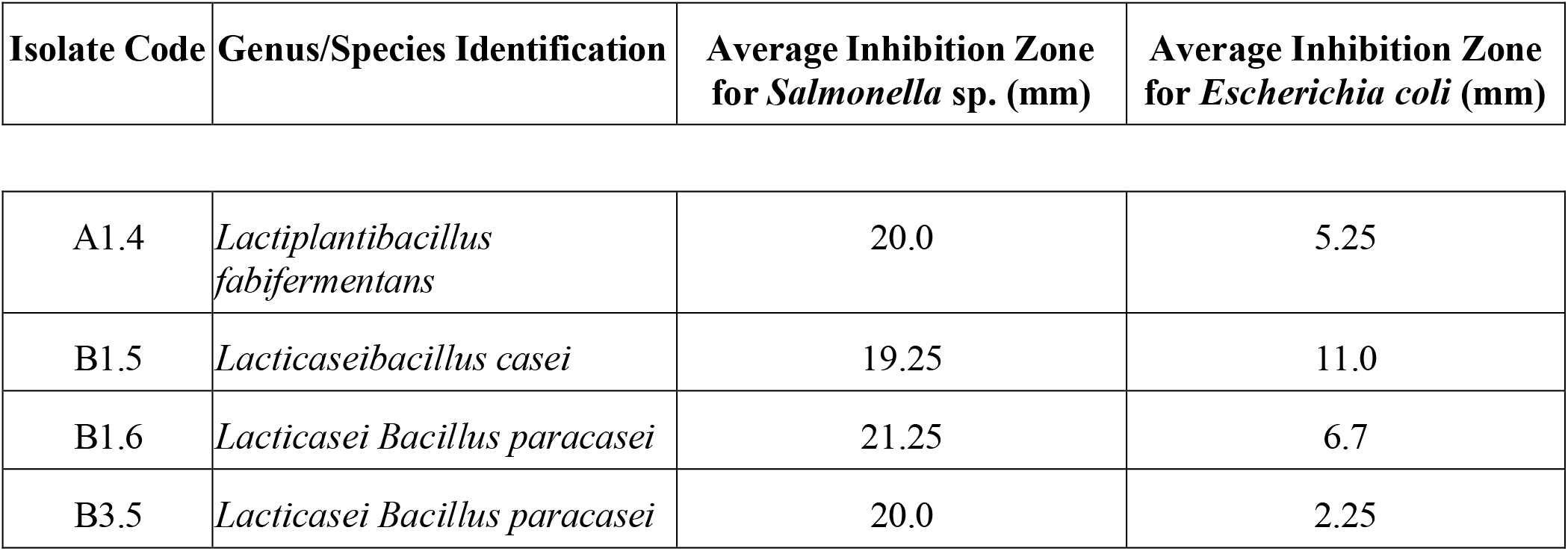
Inhibition Zone Diameters (mm) of PAL Isolates from the Wawo Palm Sugar Plantation against Pathogenic Bacteria.

| Isolate Code | Genus/Species Identification | Average Inhibition Zone for *Salmonella* sp. (mm) | Average Inhibition Zone for *Escherichia coli* (mm) |
| --- | --- | --- | --- |
| A1.4 | *Lactiplantibacillus fabifermentans* | 20.0 | 5.25 |
| B1.5 | *Lacticaseibacillus casei* | 19.25 | 11.0 |
| B1.6 | *Lacticasei Bacillus paracasei* | 21.25 | 6.7 |
| B3.5 | *Lacticasei Bacillus paracasei* | 20.0 | 2.25 |

The antibacterial activity of BAL isolates from the palm sap of the Wawo Plantation exhibited a very broad spectrum of inhibition, particularly against *Salmonella* sp. All isolates demonstrated superior inhibitory activity. The widest inhibition zone was observed in the *L. paracasei* isolate (B1.6), with a diameter of 21.25 mm, followed by isolates A1.4 and B3.5 at 20.0 mm. Even the lowest inhibition zone still showed a significant protective value of 19.25 mm (isolate B1.5).

**Figure 2.**
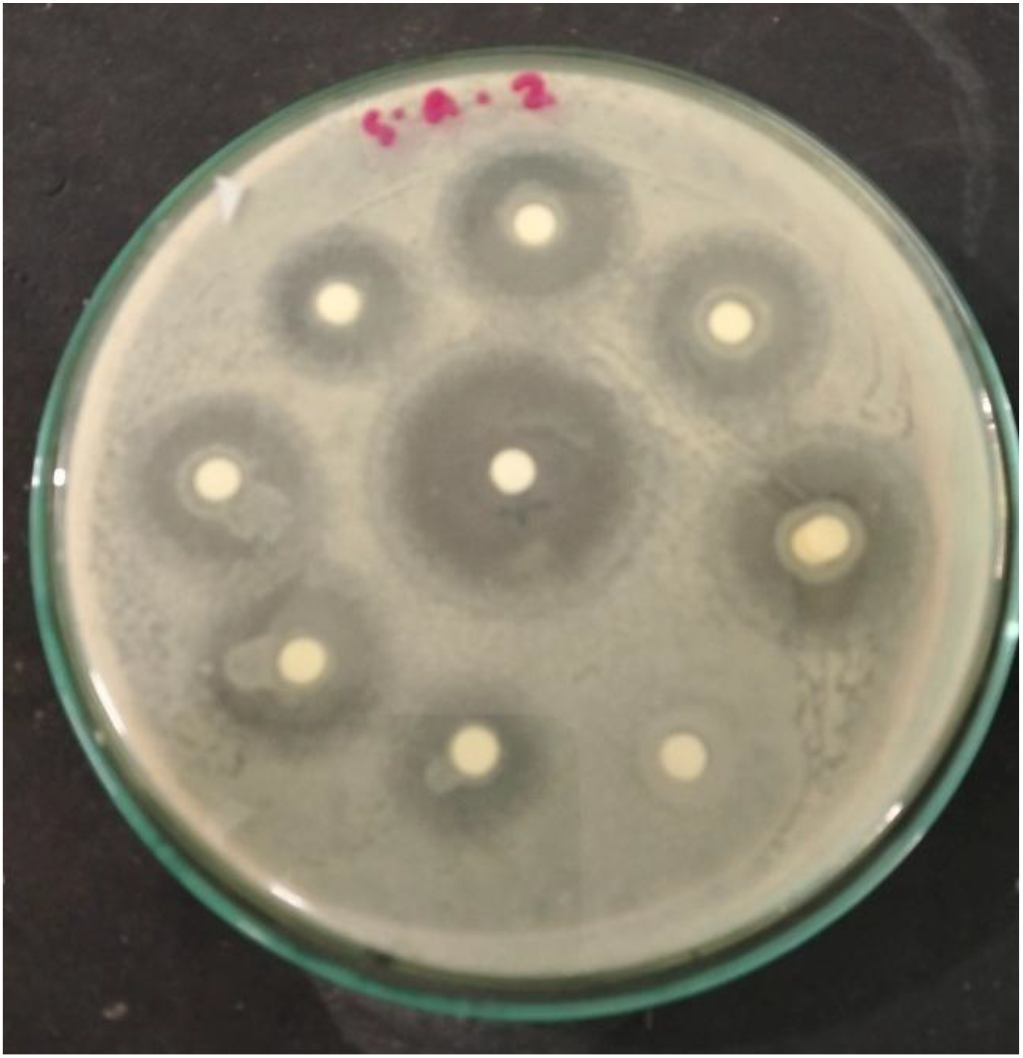
Inhibition Zones against Salmonella sp.

On the other hand, the inhibition rate against *E. coli* showed much greater variability. The *L. casei* isolate (B1.5) emerged as the most effective isolate, inhibiting *E. coli* growth to a diameter of 11 mm, followed by isolate B1.6 (6.7 mm).

**Figure 3.**
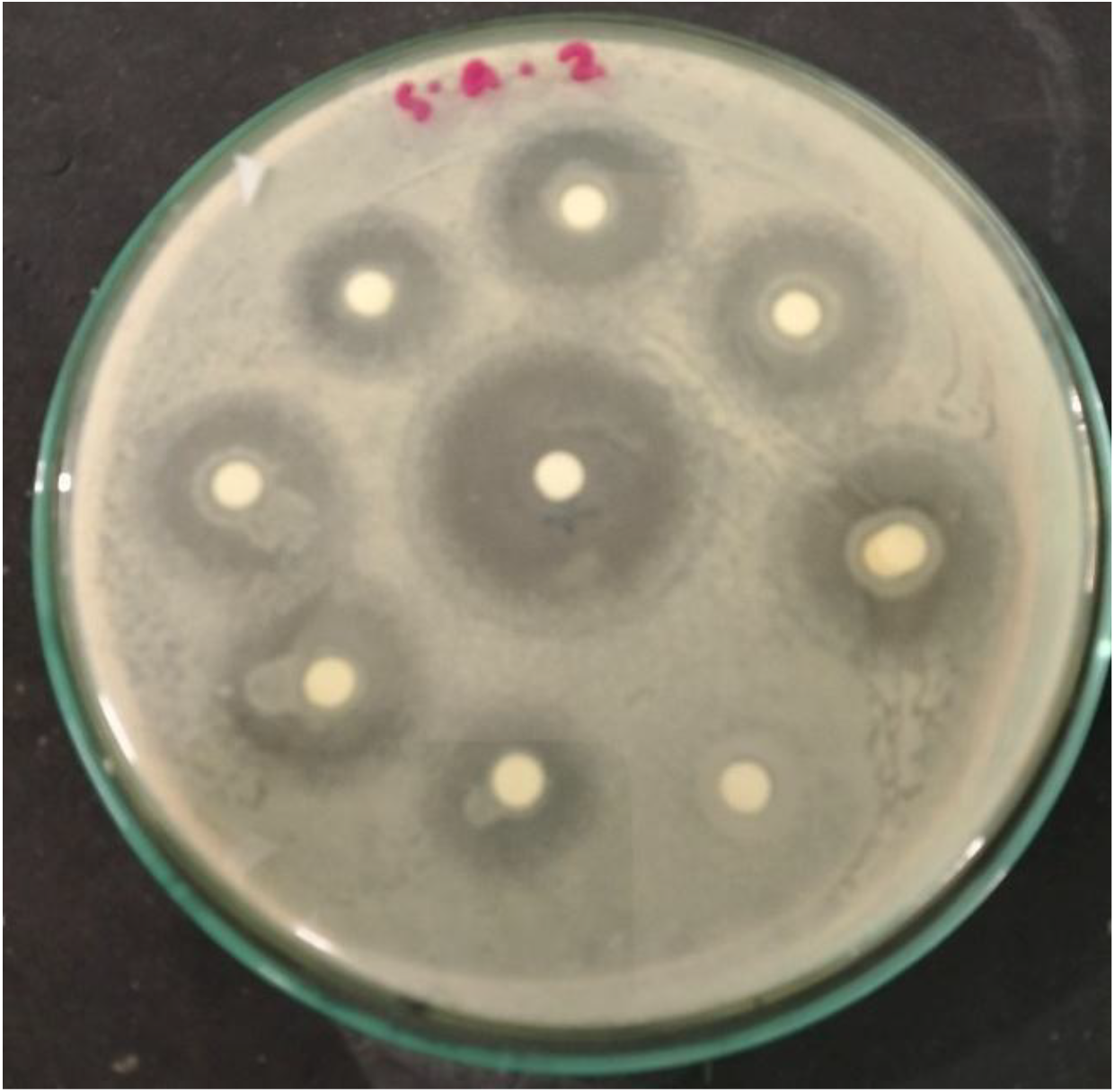
Inhibition Zones against Escherichia coli.

The antibacterial activity of the BAL Wawo strain appears to be superior and competitive when compared to similar studies in the literature. By way of comparison, in a study conducted by (Petronela, 2019), BAL isolates derived from lontar palm sap substrate were also reported to possess broad-spectrum activity in inhibiting the multiplication of *Salmonella Enteritidis* and *Escherichia coli* cells, although the width of the inhibition zone depended strictly on the level of extracellular metabolism. Furthermore, when compared to the study “regarding *Lactobacillus casei* derived from conventional fermented milk media against the pathogen *Escherichia coli*, local palm sap isolates (B1.5 and B1.6) exhibited a more dominant inhibitory interaction on the test plates.

The high inhibition zone diameters observed in isolates such as *L. paracasei* and *L. casei* Tomohon demonstrate the potent genetic mechanisms of local biotypes in producing and secreting low-molecular-weight organic acids (such as pure lactic acid), hydrogen peroxide, and specific bacteriocins. These secondary metabolites are highly active in destroying lipid components and disrupting the permeability of the outer cell membranes of Gram-negative bacteria (Asriani et al., 2007; Shavira et al., 2022). This highly effective inhibitory activity against enteropathogens confirms that lactic acid bacteria from palm sap have high potential— not only as biopreservatives in the local food industry supply chain but also as crucial candidates for future nutraceutical agents in protecting the physiological integrity of the host’s gut microbiota.

## 4. Conclusion

Lactic acid bacteria isolates that dominate the natural ecosystem of palm sap (*Arenga pinnata*) from the Wawo Plantation production center in Tomohon City have been successfully characterized and genetically identified using 16S rDNA sequencing. These isolates were identified as belonging to the bacterial groups *Lacticaseibacillus casei, Lacticaseibacillus paracasei*, and *Lactiplantibacillus fabifermentans*. The four leading isolates— —exhibited a rod-shaped morphology, were Gram-positive, non-motile, and catalase-negative. The results of *in vitro* antimicrobial susceptibility testing demonstrated highly significant broad-spectrum inhibitory activity. The *Lacticaseibacillus paracasei* (B1.6) isolate exhibited the highest inhibitory effectiveness against the spread of *Salmonella* sp. (21.25 mm), while the growth of enteropathogenic *Escherichia coli* was most strongly suppressed by the *Lacticaseibacillus casei* (B1.5) isolate, with an inhibition zone of 11.0 mm. Given its performance, which surpasses that of some BAL profiles derived from other media sources, the rich biodiversity of this superior local palm sap biotype warrants its formulation as an innovative, *food-grade* natural *biopreservative* or a nutraceutical probiotic agent.

